# End-to-end automation of behavioural experiments with a collaborative robot

**DOI:** 10.64898/2026.10.20.758951

**Authors:** Riddha Manna, Tomislav Štampar, Ivan Tomić, Ana Marija Jakšić

**Affiliations:** École Polytechnique Fédérale de Lausanne, School of Life Sciences, Brain Mind Institute, Lausanne, 1015, Switzerland; TS Robotics, Landshut, 84036, Germany; École Polytechnique Fédérale de Lausanne, School of Engineering, Discovery Learning Laboratories, Lausanne, 1015, Switzerland

## Abstract

Reliable behavioural measurement is critical to neuroscience discovery, but end-to-end automation of experiments using live animals is challenging. Preparation and transfer of live animals into and out of behavioural assays require skilled, dexterous, gentle and consistent handling. This is difficult to achieve without bespoke automation systems, which resist wide adoption. We present **STELA**, a robotic cell featuring off-the-shelf collaborative robot (cobot) that autonomously runs the complete workflow of a *Drosophila* behaviour experiment, from vial to data. Designed to emulate human routine fly laboratory procedures in human workspace, it performs manipulation of fly culture vials, CO_2_ fly anaesthesia, vision-guided sex classification, single live fly manipulation, and loading and disassembly of a behaviour assay, all in a programmed loop. In 24-h working day, STELA completed 16 experiments (1024 individual visual learning assays in flies) compared to the human who, in an 8-hour working day, completed 6 (384 assays). Retention of flies fit for analysis was similar (93.85% versus 83.33%, respectively), but with 12-fold improvement in handling consistency by the robot. In biologically matched behaviour experiments, robotic and manual handling resulted in the same mean fly behavioural outcomes, yet the robot significantly reduced inter-individual variability in baseline and learning behaviours. STELA demonstrates that it is possible to deploy standard robotic systems to flexibly automate one of the most challenging animal experiments. It provides a low-variance, high-throughput platform for scalable experimentation, and a path to closed-loop, democratized, robotic acceleration of experimental discovery.

**One-sentence summary:** A dual-arm collaborative robot emulates the full manual workflow of a fly behaviour experiment, delivering a thousand individual cognitive assays per day with lower experimental variance and higher throughput than a trained human operator.

## Introduction

Individual behaviour is the ultimate functional readout of the nervous system. Its precise and reproducible measurement is central to behavioural biology and neuroscience. Individual behaviour emerges from the animal’s integrated sensory processing of the external world, its internal state and past experiences. These factors are known to affect cognitive behaviours, including variation in learning outcomes, false memory formation, erroneous recovery of true memories, and increased anxiety^1–4^. Because of that, behavioural measurement depends not only on the behaviour assay, but also on the procedures used to prepare, transfer and recover animals from the assay. These procedures must be carefully specified and executed. The control of technical variation associated to animal handling is particularly relevant in studies of individual behaviour^1, 5–7^, where variation in handling can translate into the measured inter-individual variation. In conventional laboratories, animal selection, transfer, preparation, assay loading, and subject recovery remain manual. This creates substantial experimental labour demands as well as opportunities for inducing procedural variation between laboratories and experimenters. The effects of procedural variation on animal behaviour has been widely documented. Even the experimenter’s own biology, such as their sex, smell and identity, can significantly affect experimental outcomes^2, 8–12^. Automating the complete experimental workflow with machines or robots could offer a solution to such experimental variability as well as increase scalability.

Automation has transformed biological experimentation, especially the *in vitro* and *ex vivo* phenotyping^13–18^. *In vivo*, behavioural phenotyping has seen comparatively modest gains from process automation. For example animal pose tracking and measurement of locomotor-, feeding-, learning-, social- and other behaviours^1, 19–25^, have been individually automated allowing more scalable and quantitative behavioural analysis, but complete experiment automation has been limited. Unlike workflows in molecular- and cell biology that rely on automation of standardized liquid handling, scalable workflows that use live animals must integrate complex assay execution with delicate and safe handling of animals whose position, orientation and movement can vary unpredictably. End-to-end automation therefore poses challenges that extend beyond software-based automation and require sophisticated mechatronics. Existing end-to-end experimental automation systems that use live animals rely heavily on designing custom protocols and complex, bespoke machinery for manipulation of test subjects that must be rebuilt by the laboratory that intends to use it, and assays redesigned to fit them^22, 26–29^. As a consequence, these bottom-up automation systems are not widely adopted, despite demonstrating that they could be valuable.

Robotic automation of existing experimental protocols using a general-purpose robot could bridge this gap. Collaborative robots (cobots) offer a potential route to top-down laboratory automation by replicating the actions of a human experimenter in human-oriented workspaces, rather than redesigning the workspace and the assay to accommodate automation. Articulated collaborative manipulators combine multiple degrees of freedom with precise, repeatable positioning and power- and force-limited operation. This allows cobots to handle different tools and objects originally designed for people and to operate in workspaces shared with human experimenters^30, 31^. A dual-arm configuration further enables human-like operations involving multiple tools and objects, such as opening various containers while transferring animals between them. Organising these operations into reusable routines would allow flexible sequencing and execution of diverse experimental protocols. For behavioural experimentation, using cobots requires integration of repeatable manipulations of laboratory equipment with perception-guided handling of individual animals.

We asked can an collaborative robot be used to execute the full sequence of operations of an already established *in vivo* experiment that was designed for human operation? Importantly, can the protocol it employs and measurements it delivers remain both comparable and compatible with skilled manual experimentation routines? A general-purpose robot modelled on human-conducted experimentation could vastly extend laboratory automation to existing experimental equipment and protocols. This approach could enable more wide-spread automation of experiments involving live animals across laboratories.

Adult *Drosophila melanogaster* was shown to be a powerful and tractable model in which benefits of behavioural automation can be maximized. Its extensive behavioural repertoire, transgenic toolkit and suitability for large-scale experimentation have made it a central model for investigating the fundamental basis of behaviour^32–39^.

Here, we present a robotic cell that automates the routine handling workflow of a standard Drosophila experiment using a standard dual-arm collaborative robot. We demonstrate the robot’s capabilities by using it to automate an established operant-conditioning behavioural experiment designed for human operation^1^.

The cell performs a range of standard “flywork” routines such as vial handling, fly CO_2_ anaesthesia, machine-vision sex classification, vision-guided individual fly transfer, behavioural arena loading, assay initiation, and post-experiment recovery of individual flies. We sequenced these routines in a continuous closed-loop workflow to automate a behaviour assay. After the assay, recovered flies can be transferred to a collection vial or they can be identified and individually housed for subsequent use. The control architecture combines hardwired and adaptive robotic manipulation routines that allow unattended robotic experimentation in a workspace that can be shared with human experimenters.

We evaluate the system in a 24-hour work cycle comprising of 16 consecutive experiments and 1024 individual behaviour assays. We compare it to biologically matched experiments performed by a human experimenter to test the robot’s throughput and its effect on experimental biological outcomes.

By reproducing the sequence of manual operations performed by a skilled fly experimenter, the system converts conventional behavioural experimentation from a workflow executed by humans into a highly repeatable, high-throughput and autonomous process executed by robots. We call this system **STELA** (**S**calable cobot**T**-based **E**nd-to-end **L**aboratory **A**utomation), a platform designed to enable large-scale, high-precision discovery in *in vivo* experimental biology.

## Results

### A cobot cell for autonomous behavioural experimentation

The robotic cell integrates two main subsystems: a standard collaborative robot, which handles animal preparation for the behavioural experiment, and a computer station, operating the machine vision and behavioural platform for individually resolved cognitive behaviour assays^1^. The robot interfaces with the behavioural platform for experimental setup and then breaks down the experiment upon its end (Figure 1, Video 1, Video 2). The robot and the computer communicate through an FTP server to coordinate their actions using a file-based handshake protocol, in which flag files are exchanged to signal task status and trigger subsequent operations. An Arduino microcontroller connected to the Raspberry Pi controls two peripheral devices within the cell, the CO_2_ valves and the fly sieve.

**Figure 1.**
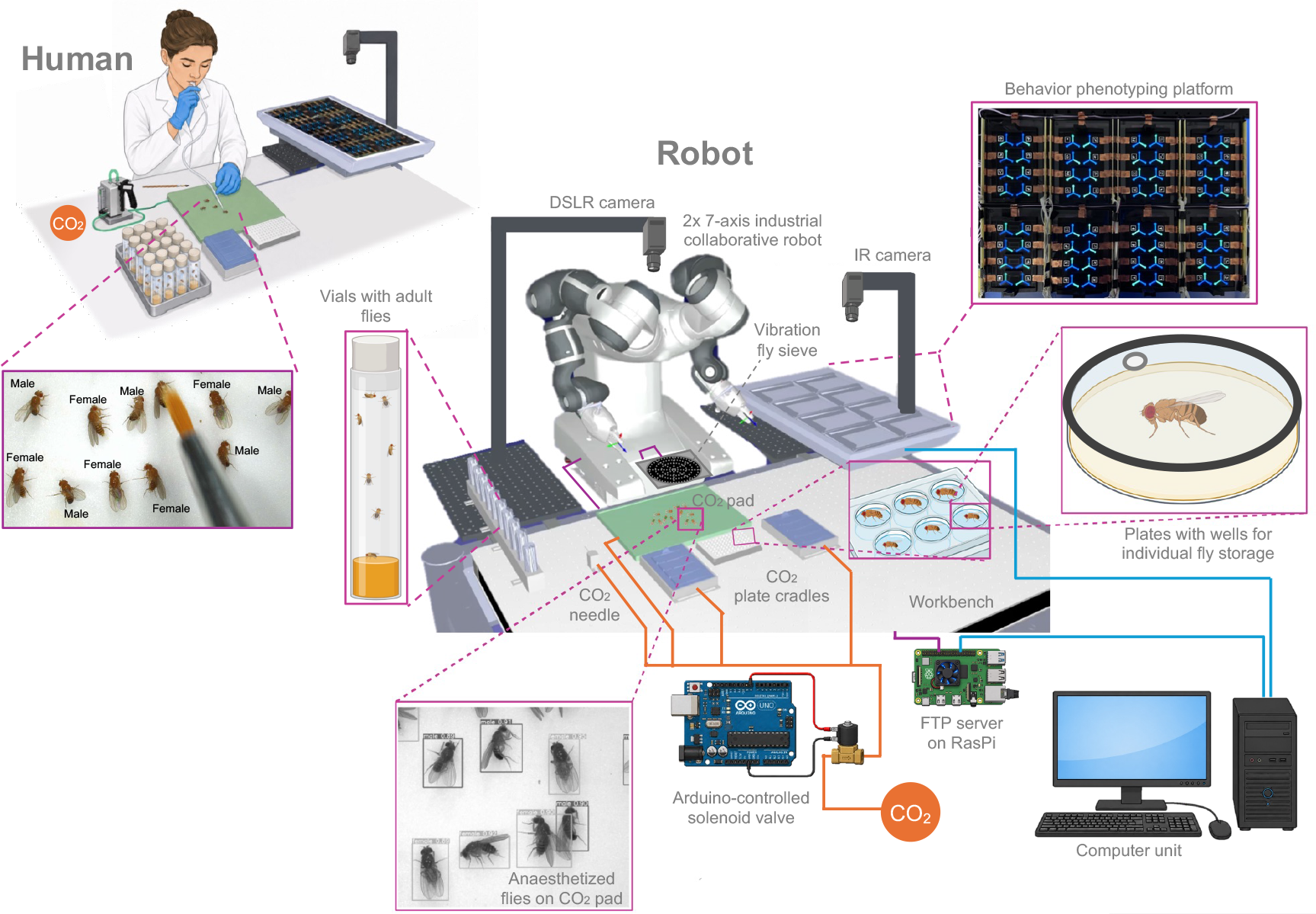
Robot cell schematic. Collaborative robotic cell is designed to emulate and reproduce human flywork in the human’s workspace, making bench top components and their organization usable by both the manual and robotic experimentation mode.

The robotic cell is built around a dual-arm, 14-degree-of-freedom collaborative robot (YuMi IRB 14000, ABB Robotics, Switzerland; Figure 2), which serves as the cell’s main actuator. The robot is equipped with interchangeable end effectors, including 3D-printed grippers, a suction cup and a vacuum-operated glass pipette for manipulating individual, *live* flies (Figure 2). These fittings are used to reproduce the operating principle of standard, handheld fly aspirators conventionally used by human experimenters for fly transfer (Figure 1). We mounted the robot on an aluminium frame rigidly affixed to a vibration-damped optical workbench (ScienceDesk with Nexus Breadboard, Thorlabs; 750 mm x 1200 mm; Figure 2). The bench top breadboard contains a regular array of threaded M6 holes, providing standardized attachment points for rapid and reproducible installation, repositioning and exchange of peripheral components. This configuration facilitates modular assembly, but the cell can as easily use other rigid surfaces to flexibly build a workspace. The cobots are designed for power- and force-limited collaborative operation with reduced operating speeds, integrated collision detection, compliant padding, and covered joints. These features permit operation side-by-side with humans, without fixed perimeter guarding or a robot cage. The cell therefore has a compact footprint and can be installed and deployed in any space equipped with a standard biology lab infrastructure such as CO_2_ supply and compressed air line for fly anaesthesia and handling.

**Figure 2.**
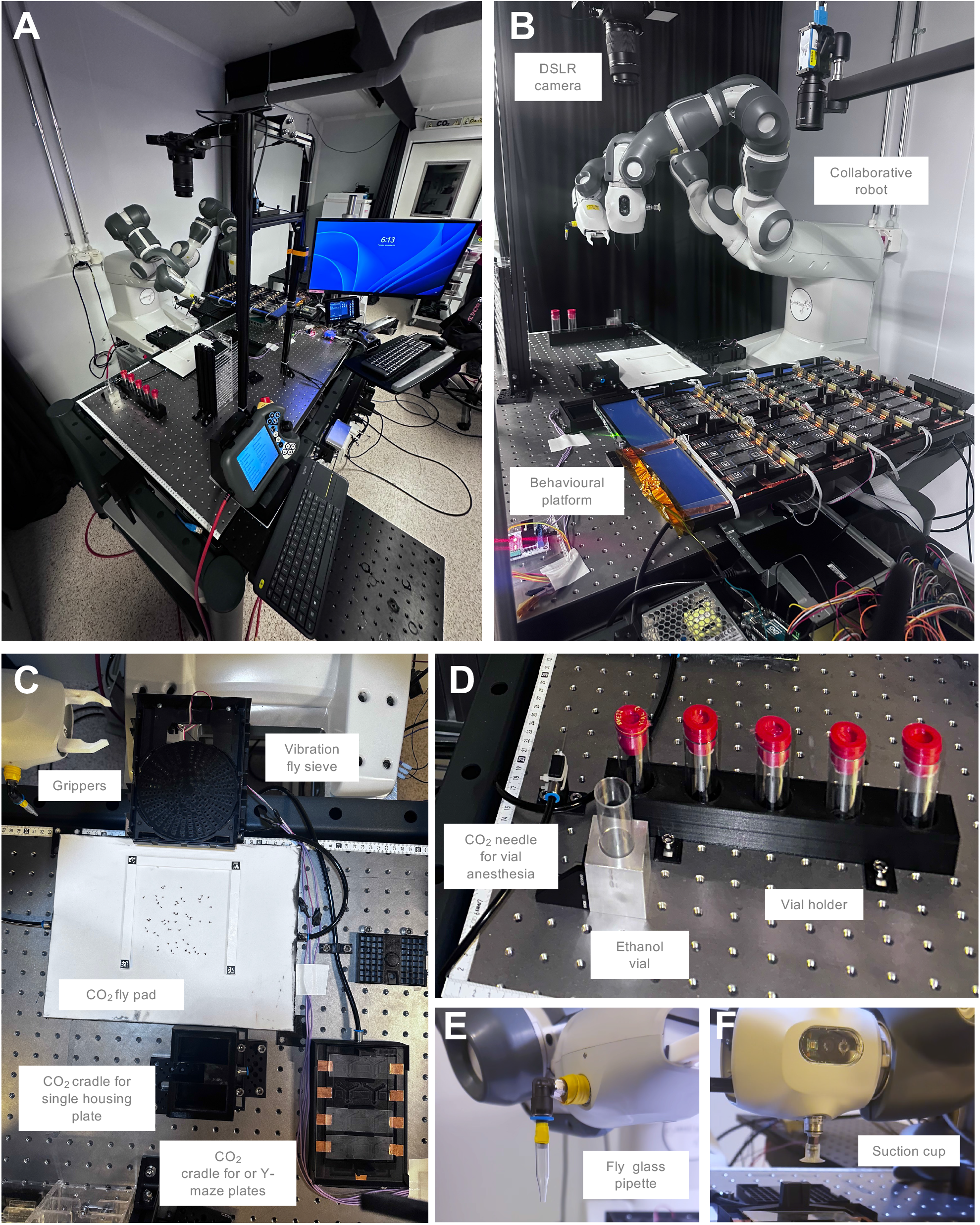
Collaborative robot cell for Drosophila phenotyping. A) Collaborative robotic cell. B) Side view of the robot (in home position) and the behavioural platform. C) Overhead view of the bench-top layout with core components for standard flywork. D) Fly vial holders and CO_2_ needle for anaesthesia. E) Custom pipette gripper for vacuum-based single fly pick-and-place. F) Suction cup for vacuum-based pick-and-place of flat items (e.g. glass slides and plastic covers).

The fixtures that interface with the robot’s end effectors are the fly culture vial holder, CO_2_ needle mount, CO_2_ anaesthesia pad, retractable vibrating fly sieve, CO_2_ cradles for Y-maze- and single-fly housing plate anaesthesia, and plate shelves. They closely replicate the standard flywork equipment and workspace arrangement (Figure 1 and 2). Consequently, the cell and all of its components remain accessible for manual operation by humans (Figure 1).

### Closed-loop control and signal-processing architecture

Robot’s actions are controlled by a combination of fixed and adaptive routines (Figure 3). The hardcoded routines handle fixed geometry and tasks that include manipulation of affixed components of the cell at known coordinates. The adaptive routines handle the irreducibly variable component of biological experimentation: the detection, classification and handling of individual flies, whose positions are randomised on each cycle. The cell is equipped with an overhead DSLR camera that acquires images of anaesthetized flies, distributed on the CO_2_ pad (Figure 2, Video 1). These images are transferred to the computer unit and processed to extract each fly’s position and sex (Figure 4). This results in a coordinate file which is then uploaded to the robot via an FTP communication server hosted on a Raspberry Pi (Model 3B). Imaging routine is triggered by the robot’s state within its action loop that signals the state to the computer via the server. Equivalently, the progression of the robot’s loop depends on the signalling from the computer when the fly position data is returned to the robot in the form of a coordinate file. This bidirectional handshake communication between the robot and the computer is mediated by flag files that are continuously uploaded, downloaded, read and deleted from the FTP server by both the robot and the computer. This makes the robot’s action loop closed rather than merely sequenced.

**Figure 3.**
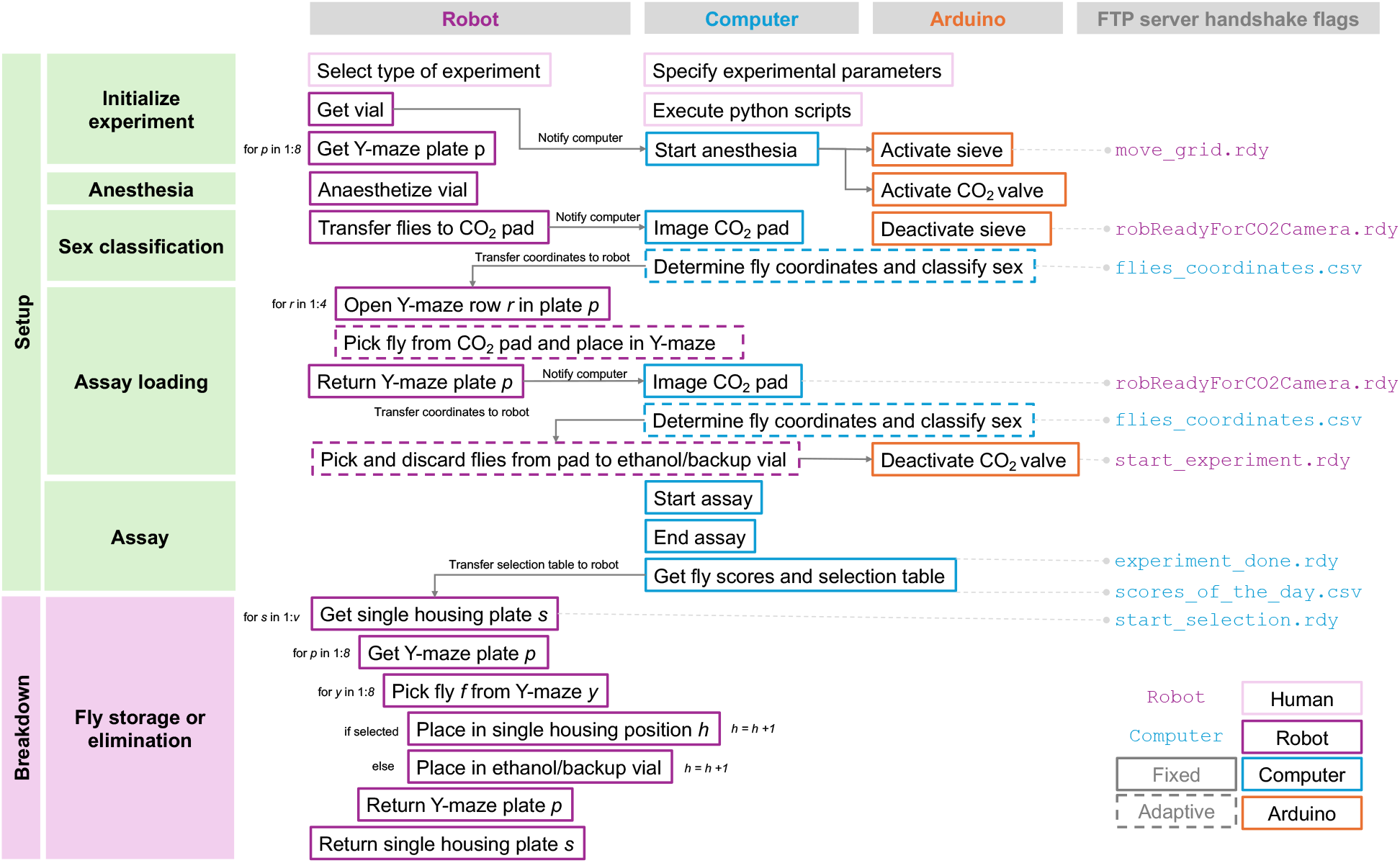
Typical example of a behaviour experiment algorithm. Standard robotic protocol for a behaviour experiment consists of different pre-programmed routines (boxes). Between each routine the robot assumes a home position. This way, the algorithm is easily broken into modules and any part of the cycle can be rearranged, added or removed via software modification.

**Figure 4.**
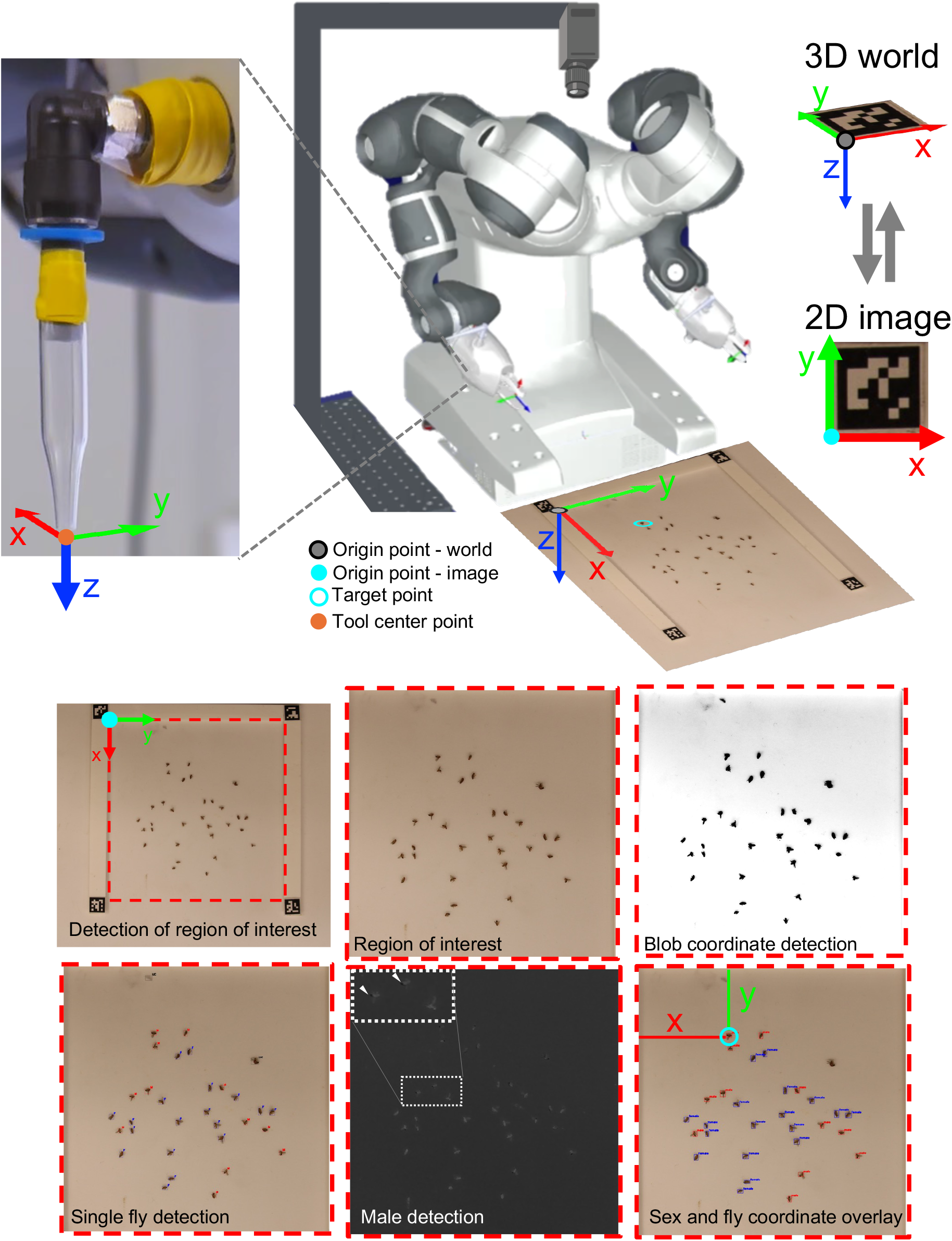
Single fly pick-and-place and sex determination. ArUco marker coordinate system is used to transform 2D fly coordinates from image to 3D world coordinates. The top left marker is used as a fixed world origin point for the CO_2_ pad. Its world coordinates are matched to pipette tip tool coordinates before every fly pickup. The imaged CO_2_ pad is cropped to a region of interest (red interrupted line), defined by ArUco markers. The cropped image is processed in steps to detect individual flies, their sex (M - male, F - female and UC - unclassified) and coordinates (target point). Inset image (grey interrupted line) zooms in on two males (arrows) and a female in the male detection step.

### Sex classification of individual unrestrained anaesthetised flies

When a human sorts and selects flies by sex (“fly sexing”), they first anaesthetise a mixed-sex vial of flies with CO_2_ gas and distribute them on the CO_2_ pad. The flies are then inspected on the pad using a stereomicroscope at 1.5–3x magnification, or by eye if the experimenter is sufficiently experienced. A brush is used to move the flies or otherwise orient them on the CO_2_ pad to observe them individually (Figure 1, Supplementary figure 1). They are then pushed on the pad to form two separated piles, one for males, the other for females. Finally, the piles are transferred from the pad into separate vials. Sex is almost always initially classified based on the melanin deposition in the cuticle of the posterior abdomen, which is much more pronounced in males than females (Supplementary figure 1). The classification is aided by the shape of the fly’s body, including genital anatomy. This procedure presents one of the most commonly required, yet at the same tedious and time-consuming manual steps in any Drosophila experimental workflow that could greatly benefit from automation. Most existing approaches to automation of fly sexing require some form of manipulation of each animal’s orientation relative to the camera, genetic manipulation, or they rely on high-magnification imaging of individuals^40, 41^. This makes these approaches complex, difficult to adopt, or slow. Here, we developed an algorithm that emulates human procedure and enables the robot to classify sex of dozens of anaesthetised flies imaged in arbitrary orientation *en masse*, and to select and handle single flies based on their sex (Figure 4).

We emulate the human process of determining fly sex using fly’s body shape and pigmentation, in several steps that employ machine vision and a simple vibrating sieve. The robot picks up a vial containing adult flies, uses a CO_2_ needle to anaesthetise them and then opens the vial over a vibrating sieve placed above the CO_2_ pad (Video 1). Vibration distributes the flies evenly on the pad as they fall through the sieve. A DSLR camera is used to acquire a single wide-field 6000 x 4000 px image of the CO_2_ pad with anaesthetised flies (Video 1, Figure 2B, Figure 2C, Figure 4). ArUco fiducial markers^42^ are used to localise a 120 x 120 mm planar workspace (region of interest, ROI) within the image and they determine the position and orientation of the ROI.

Fly detection in the region of interest then proceeds in following steps. First, a binary thresholding is applied to the grayscale image followed by blob detection filtered on area, circularity, and convexity of the blob (Figure 4). Then, a Hue-Saturation-Value (HSV) filtration is applied along with additional area and convexity criteria, which results in more confident identification of the fly based on the shape of its body (Figure 4). Overlapping flies (and smudges on the pad) fail this shape filter criteria and are ignored by the program in the following steps. Because melanin absorption is wavelength-dependent, we found it can be isolated from the photograph by subtracting the blue channel from the red channel. This transformation sharply enhances the male abdominal pigmentation signal (Figure 4, white arrows). The overlap of appropriate shape and area with the distinct posterior pigmentation signal identifies the male flies, while the rest are classified as female. Their centroid image coordinates are converted to true fly coordinates and transferred to the robot (Figure 4). In summary, as in the human procedure, robotic fly sexing is performed based on fly body morphology and pigmentation.

Overall robot’s sexing accuracy was 92.3% (N = 350 flies, 12 images), tested against the human-determined ground truth. The throughput consequence of this procedure is substantial. Manual sex classification requires 9 minutes (*±*3 min) per vial containing 70 - 120 flies (N vials = 20). This is normally performed two days before the assay takes place to avoid overexposure of flies to CO_2_ and stress-related classification mistakes by the experimenter. In the robotic cell, the same procedure occurs right before the assay. It takes around 40 s per vial, including vial handling (on average 20 s), image acquisition ( 20 s) and the classification itself (200 ms), making it an order of magnitude shorter.

### Single fly pick-and-place

To determine the true fly coordinates, one corner of the planar workspace in the image defines the origin of the local coordinate frame registered to the robot base frame. The fiducial markers provide a planar homography between image-pixel coordinates and the corresponding X-Y coordinates on the CO_2_ pad. Detected fly positions are transformed from image coordinates into robot base frame at the known Z-height of the pad. The resulting 3D target coordinates are logged and transmitted to the robot controller. The controller then generates the motion trajectories for the tool centre point (TCP) of the glass-pipette end effector during the fly pick routine (Figure 3, Figure 4).

The robot uses the coordinates to pick up and load males and females in various containers using gentle aspiration and controlled air blow-out. The containers can be the Y-maze arena, vial, wells in single-fly housing plates or any other X-Y bound place on the workbench with a boundary height as low as 0.6 mm.

### The autonomous behaviour experiment

We demonstrate the capabilities of our robotic cell by automating behavioural experiments on an apparatus originally designed, built and validated for manual use^1^. The apparatus is a behavioural platform comprising arrays of Y-maze arenas positioned above an LCD screen. The screen delivers visual stimuli to individual flies while they explore the maze. Plates holding Y-mazes are able to deliver a very mild electric shock to individual flies in order to condition their behaviour. This configuration supports individual operant conditioning assays, a standard test in behavioural neuroscience to track and quantify learning-dependent behaviours (Figure 1, Supplementary figure 2).

To perform a complete 64-fly assay on a behavioural platform the robot cycles through several standard flywork routines normally executed by a skilled experimenter (Figure 3). Each experiment requires a fly vial containing >=64 flies placed in the vial holder.

#### Experimental setup

To start the experimental cycle the experimenter selects the experiment on the robot’s teach pendant and the robot initializes the sequence of experiment setup routines (Figure 3, Video 1). During each cycle, the robot retrieves a vial from the holder with the right arm, while the left arm retrieves the first Y-maze plate from the LCD screen and places it on the CO_2_ delivery plate cradle (Video 1). The grippers hold on to a holder on the plate, just as the human would, holding it with two fingers. The flies are anaesthetised in the vial during a 10 s-long CO_2_ delivery through a needle. The needle is inserted into the vial through the custom vial cap. The two arms then coordinate to open the vial and dispense the flies through the retractable vibrating sieve onto a CO_2_ pad (Video 1). The DSLR camera images the CO_2_ pad, and vision processing proceeds. The computer compiles and sends a file to the robot with a sequence of pick-and-place coordinates that the robot needs to follow.

The right arm, fitted with a glass pipette on a suction line, picks flies individually from the CO_2_ pad. It coordinates with the left arm, which lifts the glass slide cover from the first row of two mazes using a suction cup, while the right arm deposits one fly into each maze. The loaded mazes are then closed again with the glass slide. Four such operations fill the eight Y-mazes of a plate. The plate is then picked up again by the left arm and returned to the screen (Video 1).

The next empty plate is recovered from the platform and placed on the cradle. The camera re-images the CO_2_ pad and updates the positions of remaining flies (because lightly anaesthetised flies may still slightly move), and the whole sequence repeats until all 8 plates (in total 64 mazes) are loaded with flies and all Y-maze plates are returned to the screen. In this moment, the robot signals the computer to initialize the behaviour assay. Any surplus of anaesthetised flies is, in the meantime, cleared from the pad into an ethanol vial (or placed in a backup vial with fly culture media, in case they should be kept). The robot idles during the experiment to avoid disturbing the behaviour of the flies.

#### Experimental breakdown

On completion of the assay, the robot is notified by the computer and proceeds to retrieve the plates, in what is essentially a reverse sequence of the experiment setup routine. The robot re-anaesthetises the flies on the CO_2_ plate cradle, lifts the glass cover and recovers individual flies from the Y-maze by hovering the glass pipette in a Y-shaped motion over the maze, while keeping a sustained suction (Video 2). The recovered flies are either discarded in ethanol, stored in the backup vial, or transferred to a single-housing 64-well plate for downstream use in which fly identity needs to be preserved (Video 2). The empty Y-maze plate is returned to the screen and the sequence re-iterates until the entire platform is unloaded.

If flies are unloaded into a single-housing plate, at the end of each experiment cycle the single-housing 64-well plate is retrieved from the plate shelf, filled with flies and then stored back on the shelf so that it can be accessed by the robot or the experimenter at a later time point. Each plate is stored on a numbered shelf, with the number matching the experiment cycle number.

#### Post-cycle options

One complete cycle (setup, 25 minute assay and breakdown) takes approximately 75 min (Figure 5A, 5B). If flies are stored individually, the scores_of_the_day.csv file can be used to select flies of interest based on their behaviour scores computed from the assay data at the end of each assay. Selection can be done at the end of each cycle or at the end of each day, using the same single fly pick-and-place routines. The flies can therefore be selected straight from the Y-maze or from the wells in the plate (Video 2). After six experiment cycles or once the provided vials of flies have been exhausted, the robot prompts the experimenter for vial replacement via the robot’s teach pendant, via email, or a phone message. The limit of six cycles was initially arbitrarily set by the first vial holder design but can be easily extended by redesigning the vial holder and via standard software modifications to robot arm trajectories.

**Figure 5.**
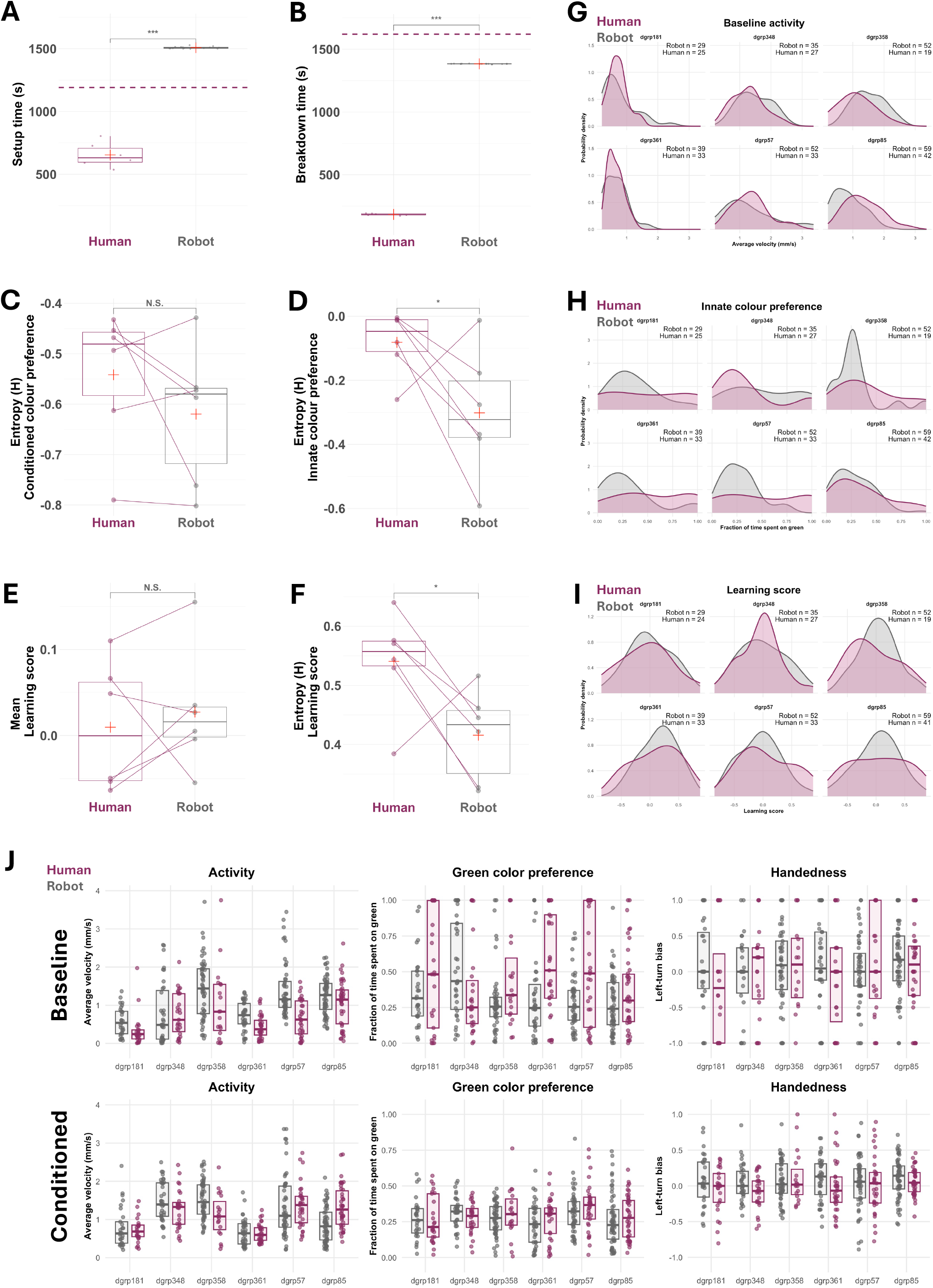
Robot versus human performance. **A)** Setup and **B)** breakdown time for 64-fly behaviour experiments. Each point is an experiment (human N=6, robot N=16). Dashed magenta lines show estimated average time spent on manually setting up the experiment including fly sexing done 2 days prior to the experiment (mean experiment setup time + mean fly sexing time, 1192 s), and average time spent on manually dismantling the setup when individual fly identities were preserved (mean breakdown time from multi-day experiments in a previous study^1^, 1620 s). Expressed variability (entropy) of **C)** conditioned colour preference and **D)** baseline/innate colour preference in six genotypes (points). **E)** Mean and **F)** expressed variability (entropy) of learning score in six genotypes (points). Inter-individual variability is significantly lower for innate colour preference and learning, but not for mean learning score. **G)** Genotype-wise distributions of individual baseline activity show similar spread between robotically and manually handled flies. **H)** Genotype-wise distributions of individual innate colour preference show higher inter-individual variability for manually handled flies. **I)** Genotype-wise distributions of individual learning score show higher spread for manually handled flies. **J)** Differences in expressed individual behaviours between robot- and human-handled flies are most prominent in baseline measurement of behaviours, which are measured right after handling (each point is an individual fly; boxes show median and inter-quartile range).

We successfully deployed the main loop architecture and here we report results from 19 cycles (Video 1), including one 24-h continuous run of 16 consecutive cycles that we use to validate the robot’s performance against the human’s (Supplementary table 1). This demonstrates that the robotic cell can successfully automate an experiment initially designed for human experimenters.

### Performance against a skilled human experimenter

We used two types of experiments, *A.) mutant behaviour experiment* and *B.) wild-type behaviour experiment*, to test the performance of the robot against the human (Supplementary Table 1). In both experiments, we use the same operant conditioning assay implemented on the behaviour platform. Cycle time performance was tested using the data of the complete wild-type behaviour experiment. Differences in biological outcomes were tested in mutant- and the wild-type behaviour experiments, but on the subset of genotypes that were handled both by human and the robot (Supplementary Table 1). We measured the time performance in setting up and breaking down a full behaviour assay in wild-type behaviour experiments. We benchmarked the robot’s performance, where possible, against the performance of a highly experienced human experimenter (who has previously set up >100 such experiments) executing an identical experiment. We also measured the consistency in biological outcomes between experiments set up by both human and the robot. In the experimental breakdown the recovered flies were discarded after the assay.

#### Cycle time

Setup of a full 64-fly assay cycle took on average 1509 *±* 8 s for the robot (mean *±*s.d.; Figure 5A) versus 1192 s manually (including 654 *±*97 s loading the plates and 538 *±*166 s for sex separation 2 days prior to the assay; Figure 5A). Manual sex separation is performed before the assay to allow flies to recover from longer CO_2_ exposure. The difference reflects primarily the tool centre point speed limit of a collaborative robot which was kept at 500 mm/s to guarantee safe handling of glass components and live animals.

The breakdown of the assay took 1384 *±* 2 s (Figure 5B). The breakdown part of the cycle could not be directly compared against the human because the procedure was not kept the same. In manual unloading of a Y-maze plate the fly identities are not tracked: the plate is placed on a CO_2_ pad to anaesthetise the flies and all four glass covers are removed at once. Then, arenas are unloaded all at once, by tipping the entire plate and dropping all 8 flies on the CO_2_ pad. The flies are then vacuumed from the plate with a hand-held vacuum cleaner. This procedure took 184 *±* 8 s (Figure 5B). The robot, on the other hand, always processes arenas and flies serially and gently recovers flies one by one from each maze, regardless whether they are to be discarded, stored in groups or individually. While the manual technique is substantially faster, the serial unloading allows post-assay recovery of flies with resolved identities and even the flies that will be discarded are handled gently. Although this approach is also technically feasible with manual handling, it is rarely used. From five such manual experiments previously done in our lab that implemented serial unloading (reported as “multiday experiments” in^1^), we approximated that serial, manual, identity preserving experiment breakdown takes on average 1620 s, an order of magnitude longer than the usual tipping method we report here. Accurate preservation of fly identities requires much higher concentration and dexterity from the experimenter than Y-maze loading. It is arduous, repetitive, and physically and cognitively fatiguing, making this procedure generally undesirable from occupational-welfare perspective. This is why we avoid it whenever possible, including in this study. From experience, increased operator fatigue also greatly increases the risk of fly injury or loss through mishandling, as well as errors in maintaining individual fly identities.

#### Repeatability

The same robot cell hardware configuration, control software, and operating procedures should facilitate replication across laboratories, and can thus provide experimental standardization as well as circumvent biases dependent on experimenter’s own biology^2, 8–12^. In support of this, the robotic cell exhibited substantially greater temporal consistency in handling flies than the human experimenter (Supplementary figure 3). The standard deviation of the experiment setup time was 8 seconds for the robot (N=16) against an order of magnitude larger deviation of 97 s for the human (N=6; Figure 5A). For experiment breakdown, the corresponding standard deviations were 2 s and 8 s, respectively, despite the robot’s procedure comprising of a much longer sequence of serial single fly recovery (Figure 5B, Video 2). These values represent approximately 12-fold and 4-fold reductions in technical variability, of setup and breakdown times, respectively. The greater variability of human operation was, from experience, primarily associated with handling interruptions, such as dropping a brush or a fly, and time-dependent operator fatigue (Supplementary figure 3), while the robot’s variability depended primarily on the spatial distribution of the flies as they land on the CO_2_ pad. These results demonstrate that robotic automation can reduce cross-experiment variability caused by inconsistent handling times as well as remove the dependence of experimental consistency on the performance of a single person.

#### Throughput

Over the 24 h period, the robotic cell could complete 16 experiment cycles, yielding 1024 individual learning behaviour measurements. Operation during this time was unattended, except mid-experiment vial replacement, which did not require a skilled operator. By comparison, during the same work period, a skilled experimenter could perform up to 6 cycles (provided that the serial unloading procedure was omitted). This sums up to a maximum of 384 tested flies within an 8-h working day that included breaks to rest, and with the entire working period dedicated solely to this workflow. The robotic cell therefore achieved at least 2.7-fold greater daily throughput than a single human experimenter. Automation additionally released the skilled experimenter from repetitive and physically demanding experimental tasks, allowing their time to be redirected towards cognitively more stimulating, creative scientific work.

#### Setup quality and loading errors

Flies’ condition after being loaded into the maze was first assessed using lenient retention criteria to identify individuals that remained sufficiently active and fit for measurement of any behaviour. Similar proportions of robot-handled flies (93.85%, n/N=961/1024) and manually handled flies (83.33%, n/N=280/336) were retained. Behavioural analyses used additional, stricter behavioural filtering criteria to exclude unfit flies and flies for which it was not possible to measure all behaviour scores (see Methods, Table Y). After behavioural filtering, 74.32% (n/N=761/1024) of robot-handled flies and 53.27% (n/N=179/336) of manually handled flies remained for learning- and other behavioural analyses. During robotic handling, 8 mazes were left empty and 3 contained more than one fly. Review of fly recordings suggested that some robotic loading errors arose from missed or double pickups from the CO_2_ pad. The errors have also been caused by flies adhering to the pipette walls or opening, and being displaced unpredictably by airflow during deposition (Supplementary video 1).

#### Preservation of biological outcomes

Automation of behaviour experiments is useful only if it preserves the animal’s normal behavioural responses. We tested three genetic lines of flies: the blind mutant line (*norpA^EE^*^5^), a memory-deficient mutant (*dnc^1^*), and a wild-type genotype (DGRP 362) that we previously showed to have a pronounced visual learning ability^1^. We tested whether the robotic handling affected learning behaviour in freely moving flies. In this assay, individual flies explored a Y-maze in which one arm was illuminated in green and the other two in blue colour. During *conditioning*, entry into the green arm triggered a mild foot shock, while exiting green removed the shock, thus training the fly to learn a task of avoiding that colour. We measured task performance (green colour preference) as fraction of time spent on green or shocked arm (T_shocked_), where a decrease in T_shocked_ from 0.33 is indicative of successful task performance (Supplementary figure 2B). Before conditioning, each fly explored the arena without reinforcement or punishment, providing a *baseline* measure of innate, spontaneously expressed behaviours, including colour preference. Learning was measured as a change in behaviour from that baseline after conditioning the flies with the shock (Supplementary figure 2A). Each genotype was also tested in a *control* experiment where no shock was ever applied, and thus no learning to avoid green colour was expected.

After being loaded in the assay by the robot, the conditioned wild-type flies demonstrated a significantly better task performance (T_shocked_ = 0.17 - lower is better) compared to the control flies (T_shocked_ =0.35, t-test P = 2.47e-04, N = 64), as expected. The blind (conditioned T_shocked_ = 0.25, control T_shocked_ = 0.29, t-test P = 0.19, N = 64) and memory-deficient (conditioned T_shocked_ = 0.28, control T_shocked_ = 0.27, t-test P = 0.75, N = 64) mutant flies, did not learn to avoid the green arm, as expected due to their genetic sensory and cognitive impairment. These results replicated the results obtained after manual setup (Supplementary figure 2B). We found no significant difference in task performance between flies handled manually and those handled by the robot in any of the experimental groups. This indicates that the that robotic handling preserves the normal average expression of both the wild-type and mutant colour learning behaviours. The two modes of handling, robotic and manual, are thus experimentally interchangable when comparing mean behaviour across different groups (Supplementary figure 2B).

### Robotic handling yields lower inter-individual variability while preserving measurement means

Robotic handling could induce fine differences in behaviour compared to manual handling. Because we used mutant genetic lines expressing extreme phenotypes, those fine differences could be obfuscated when comparing group means. We therefore tasked the robot to set up 6 experiments, measuring 384 flies in 6 randomly selected wild-type genotypes. An experimenter also manually set up six identical experiments, using genetically identical flies raised in the same environmental conditions. This ensured that handling mode remained the only experimental variable. We measured a range of individual behaviours expressed throughout the assay, including locomotor activity, colour preference, and turn bias (Figure 5J). Their baseline expression was quantified in the absence of shock while the temporal changes in behaviours measured during conditioning were used to quantify the flies ability to learn.

Mean learning phenotypes did not differ significantly between handling modes (mean Conditioned colour preference: robot handling = 0.28, human handling = 0.29, t-test P = 0.49; mean Conditioned learning score: robot handling = 0.03, human handling = 0.01, t-test P = 0.68, Figure 5E, 5I). Locomotor activity (fly’s average velocity) was also comparable between manually and robotically handled flies, both during the baseline measurement period (robot handling = 1.03 versus human handling = 0.71 mm s^-1^, t-test P = 0.14) and conditioning period (robot handling = 1.11 versus human handling = 1.07 mm s^-1^, t-test P = 0.86; N = 445 flies).

We next examined whether robotic handling affected inter-individual behavioural variability. This is a particularly important measure when studying individual behaviour. In such studies, biological variation must be separated from technical variation caused by individually experienced conditions, for example variation in handling of individual animals. Variability across individuals within a group of flies was quantified using an entropy-based measurement of variation, as previously described in^1^. Innate colour preference entropy was significantly lower among robotically handled flies (mean H = −0.30) than among manually handled flies (mean H = −0.08; t-test P= 0.04; Figure 5D), consistent with robot’s reduced handling-related variability during baseline measurements. On the other hand, the entropy of conditioned colour preference did not differ between handling modes (Figure 5C). This reduction is consequential because individual learning behaviour is quantified relative to pre-conditioning behaviour. Variability in the baseline therefore propagates directly into individual learning metrics. Consistent with this, we find broader spread of individual learning scores in flies handled manually, compared to those handled by the robot (Figure 5F, 5I).

Because we found no significant difference in behaviour means between the handling systems, the excess in inter-individual variability observed in manually handled flies likely arises from stochastic inconsistencies in manual handling of individual flies. This would also suggest that the robotic handling essentially emulates the average manual handling of individual flies that is highly repeatable.

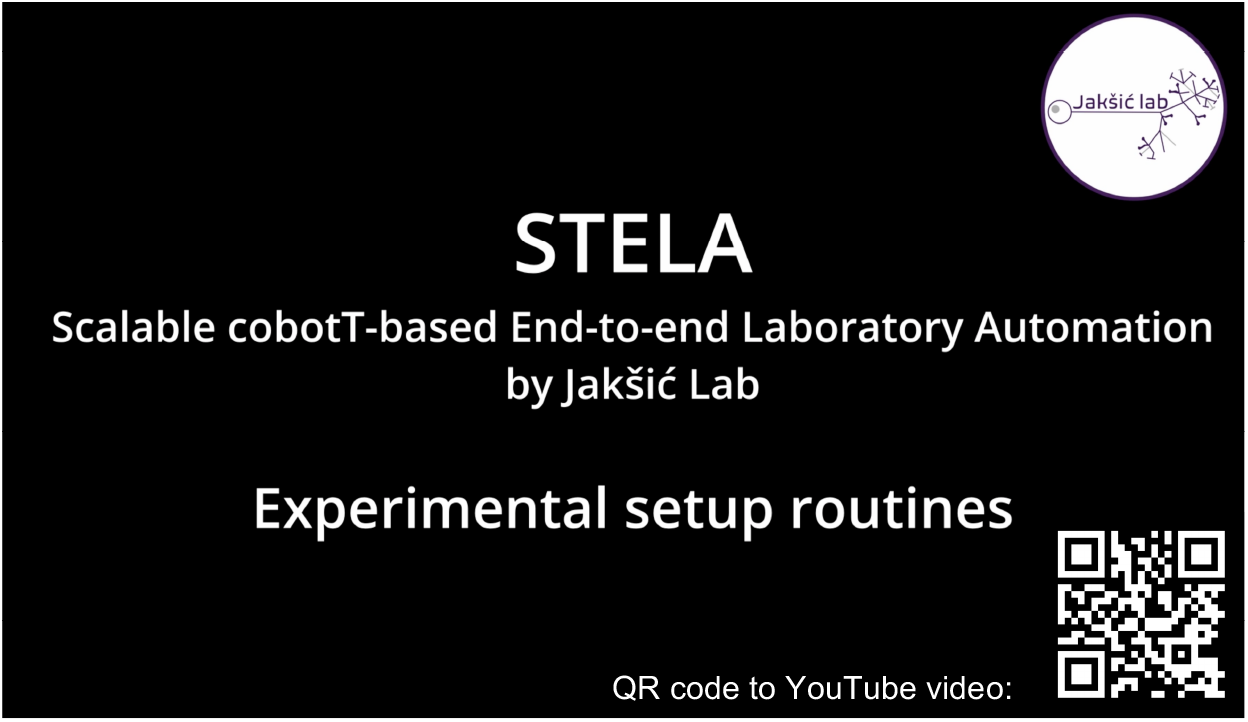

**Video 1. Experiment setup.** A dry cycle run of an example experimental setup sequence used to produce behaviour data in this study. The video on the right side shows the full cycle sequence during a training dry cycle without flies. In training trials, non-glass components of the cell are used to test and optimizing the sequence before deployment. This avoids breaking often precious glass components in case of mis-programming the trajectories while implementing a new sequence of routines. Video on the left side provides a close-up recording of each part of the cycle filmed during several real experiments. Note the consistency in movement between the trial dry run and any real experiment.

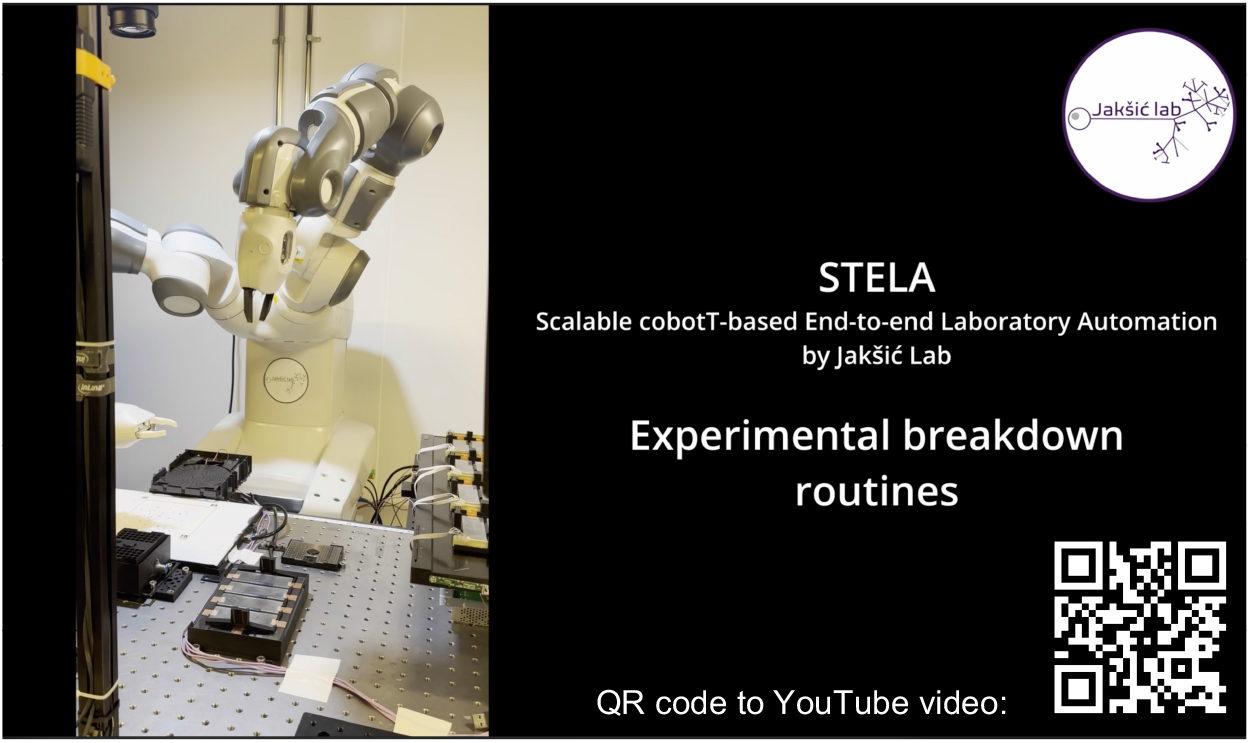

**Video 2. Experiment breakdown routines.** Recording of example experiment breakdown routines, including single-fly storage in a shelved 64-well single-housing plate or group storage or discarding of flies in a vial. The 64-well plate can be placed on fly culture media for multi-day storage and re-testing of flies.

## Discussion

Our results demonstrate that general-purpose collaborative robots can serve as adaptable and highly consistent laboratory operators, capable of executing complex experimental workflows involving live animals. STELA extends this approach to one of the most challenging biological domains to robotize: quantitative behavioural experimentation. Rather than developing bespoke mechatronic solutions for individual assays, STELA translates established manual laboratory procedures into programmable robotic operations executed by an off-the-shelf collaborative robot^26, 27, 29, 43^. By using a standard cobot, we combine the flexibility of human-operated workflows with the precision, repeatability, and autonomy of robotic execution. This approach also lowers barriers to adoption and democratizes robotic experimentation by making it more accessible to laboratories without specialized mechatronic engineering expertise or access to manufacturing facilities. We demonstrated that STELA autonomously executes the complete experimental workflow, from manipulation and preparation of delicate, live animals to behavioural data acquisition, while maintaining experimental quality comparable to that of skilled human operators.

Beyond sustained 24-hour operation and increased experimental throughput, a central advantage of STELA is the reduction of operator-dependent experimental variability. Robotic execution exhibited approximately twelvefold greater consistency in experiment setup timing than manual operation by a trained and experienced experimenter. Importantly, our behavioural entropy analysis revealed significantly reduced variability in measured baseline behaviours among robot-handled flies, consistent with a reduction in handling-associated experimental noise.

This improvement is particularly consequential for the quantitative study of learning and cognition for two reasons. First, learning is commonly quantified as a change relative to baseline behaviour, making its measurement sensitive to variation introduced before and during experimental conditioning. Second, learning can amplify initial behavioural differences over time^1^. Together, these properties may explain why cognitive phenotypes tend to be particularly noisy and susceptible to experimental variability^1–3^. By standardizing physical manipulations and their timing, STELA reduces the contribution of operator-dependent variation to measured animal behaviour. Through their own biology, human experimenters inevitably introduce differences in manipulation, timing, and interactions with animals that are difficult to eliminate entirely through protocol standardization alone^2, 8–12^. Reducing these sources of technical variation using robotic experimenters may prove to be particularly important when investigating individual behavioural differences, for which reproducibility across experiments, laboratories, and time remains a major challenge^8, 44, 45^.

STELA substantially increases the scale of behavioural experimentation. This advantage arises primarily from its ability to sustain autonomous operation over extended periods, rather than from accelerating individual manipulations alone. By reducing the need for continuous human intervention, the system also reallocates researcher time from repetitive manual procedures toward intellectually stimulating and cognitively rewarding aspects of research. Together, these capabilities open opportunities for systematic characterization of cognitive phenotypes across substantially larger numbers of individuals, genotypes, and experimental conditions, enabling behavioural studies at scales that would be difficult to achieve through manual experimentation.

Although STELA establishes the feasibility of cobot-based, end-to-end behavioural experimentation, several limitations remain in its current implementation. Arena loading errors occur in 1.1% of trials, although these can be reliably identified and excluded through video-based quality control. Automated sex classification achieves 92.3% accuracy, which remains insufficient for experiments requiring reliable sex identification at the individual level. Accuracy could be improved through repeated or independent classification, refined imaging, or improved classification algorithms, potentially at the expense of throughput. The effects of repeated robotic handling on animal survival and fitness over multiple days also remain to be systematically evaluated. Further improvements to the system are possible through optimization of the experimental cycle and robotic workspace. Currently, the robot remains idle while behavioural assays are running, and fresh animal vials require manual replenishment. Integrating a second behavioural platform would allow the robot to prepare one experiment while another is running, potentially increasing throughput substantially. Similarly, expanding vial storage capacity could extend the duration of uninterrupted operation. Following this, a priority for further development is the integration of barcode-based instead of positional vial and plate identification. This would also enable fully autonomous vial management that will eliminate the need for manual replenishment. Finally, automation of different types of assays in the future will test the limits of the robot cell’s flexibility.

Looking further into the future, STELA provides a foundation for integrating robotic experimentation with AI-driven behavioural analysis, hypothesis generation, and adaptive experimental design. The system already executes standardized experimental workflows and generates large volumes of structured behavioural data. This provides a natural interface for computational models capable of identifying behavioural patterns, detecting subtle phenotypes, and selecting experimental conditions that maximize information gain. Connecting AI-analysis and experiment design to robotic execution would enable an iterative experimental process in which results from previous experiments directly inform the design and execution of subsequent ones. Such integration could fundamentally extend the role of laboratory robotics beyond executing predefined protocols, transforming automated experimental platforms into adaptive robotic experimenters capable of navigating complex experimental spaces, refining hypotheses, and accelerating discovery in behavioural neuroscience and beyond.

## Online Methods

### Robotic cell

#### Robot

The collaborative robot used in the cell is the two-arm 14-axis ABB YuMi (IRB 14000). The robot is mounted on an aluminium stand, rigidly affixed to the optic workbench at the level of the workbench surface. The robot runs on RobotWare controller software (version 6), with programmed speeds from 30 to 500 mm/s and default acceleration settings (maximum Tool Center Point (TCP) acceleration of 39.1 m/s^2^ at nominal load). Maximum speed was limited to 500 mm/s to conform more stringently to collaborative standards.

#### Gripper fingers

Custom griper fingers were designed for the mobile components, such as fly vials, vial stoppers, Y-maze plates and single-fly 64-well plates. The right arm fingers have an indentation that match the vial circumference, and slanted finger tips that fit to gripping indentation of the plates. The left arm fingers do not have the vial-fitting indentation, and are otherwise the same. The 3D finger model was sliced with Cura, using 0.12 mm line height, 100% infill option and 3D printed on Ultimaker 5S with Ultimaker PLA matte filament. The fingers were mounted on the standard YuMi servo gripper.

#### Vacuum grippers

The left arm’s vacuum inlet was fitted with the standard YuMi gripper suction cup. The right arm vacuum inlet was replaced with a 90 degree pneumatic tube fitting with a release collet (OD 6 mm). The proximal part of the tube is affixed to the vacuum fitting perpendicular to the gripper axis, and the distal part is parallel to it. Main suction tool component is the glass pipette made from a glass 5 mL Pasteur pipette tip that was broken on two sides such that the wider, proximal side fits the release collet of the pneumatic tube, while the distal side’s inner diameter tapers to 3 mm. The sharp glass edges must be sanded. The proximal end of the glass pipette is then lined with a fine filter mesh to avoid fly injury and avoid particles entering the tube. The glass pipette is affixed rigidly to the tube fitting using insulation tape. In trial cycles the glass pipette is replaced with a simple plastic tube to avoid glass braking in case of mistakes while optimizing trajectories. Supply air pressure of 2.5 bar to YuMi vacuum module is maintained with Thorlabs air compressor (PTA524).

#### Retractable vibrating sieve

At the robot’s base, a repurposed CD-ROM drive extends a drawer over the workbench when activated and retracts it when inactive. A 3D-printed PLA disk, 120 mm in diameter and 2 mm thick, sits in the drawer and serves as a spreading surface for anaesthetised flies. Its rectangular apertures, each 3 x 2 mm, help distribute the flies evenly. The 3D model of the disk was sliced in Cura at a 0.12 mm layer height and 100% infill, and printed on an Ultimaker 5S using Ultimaker matte PLA filament. A vibration motor (Seeed Studio Grove, 9000 rpm, 5 V) is glued to the underside of the disk to help spread the flies after they fall onto it. The Raspberry Pi controls both the motor and the CD-ROM drive, automating vibration and retraction.

#### CO_2_ needle

A 3D-printed PLA hollow block with one bore and screw holders is fixed to the workbench with M6 screws. On one side of the bore a pneumatic tube fitting with a collet is inserted and sealed with silicone and can receive a CO2 inlet tube. On the other side of the bore, an intravenous catheter puncture needle (B. Braun Sterican disposable cannula, 25 G, 0.50 *→* 25 mm) is affixed and the bore is sealed using silicone. Two zip ties are used to further secure the needle in the fitting for safety, in case the seal fails under CO2 pressure. The CO2 flow in the needle is open via a solenoid valve controlled by the Raspberry Pi via an Arduino.

#### CO_2_ pad

A CO_2_ pad is designed as a slightly larger version of a standard fly pad. We printed a 300 *→* 200 mm 3D-printed PLA container with open top. A 2.5 mm thick porous ultra-high-molecular-weight (UHMW) polyethylene sheet was glued onto the top of the container (Scientific Laboratory Supplies, Flystuff (46-100) Porous Plastic Sheet) and enabled even CO_2_ dispersion from the box to surface. CO_2_ is delivered to the pad through two lateral bores fitted with a pneumatic fitting for a tube. A 120 *→* 120 mm region bounded by three 4 mm raised printed walls is affixed to the centre of the pad. defines the loading area registered in the robot firmware. One side is omitted to allow the experimenter to easily access and move the flies from the bound area if needed. CO_2_ flow is maintained at 10L/min measured and regulated with FlyStuff FlowBuddy system.

#### DSLR camera

The overhead camera Canon EOS 2000D is used for imaging the CO_2_ pad. It acquires a 6000 x 4000 px image of the CO_2_ pad with 70 mm lens and following parameters: f/8, ISO 100, 0.25 s). The camera is mounted on aluminium profiles 1000 mm above the bench top. The focus was manually adjusted and locked at the level of the CO_2_ pad, such that the flies are in sharpest focus.

#### CO_2_ delivery cradles for Y-maze plate and single-fly 64-well plate

A hollow 3D-printed PLA holder (160 x 120 x 30 mm), connected to the CO_2_ supply via a pneumatic valve, holds the Y-maze plates and delivers CO_2_ to anaesthetise the flies in the Y-maze during setup and breakdown of an experiment. The edges of the holder are slanted to accommodate small misalignments between the Y-maze plate and the holder if gripping the plate was imprecise. Similarly the cradle for 64-well plate is designed the same way, but its geometry is U-shaped and the holder is elevated from the bench top to allow sideways gripping.

#### Vial holder

The vial holder used in this setup is a 3D-printed block measuring 140 mm x 45 mm with five vertical bores, closed on bottom side. Each bore diameter is 26 mm in diameter, designed to house five standard 25 mm diameter fly vials and can be used for manual fly work.

#### Vial cap

The custom vial cap is designed as a single 3D-printable part in flexible thermoplastic polyurethane (TPU) for standard 25 mm fly culture vials. Its conical lower profile eases insertion and removal, while graded infill (0% at the base to 100% at the top) provides a compliant base for sealing and a rigid top for robotic gripping (Supplementary Figure 5). The hollow upper section has an integrated 0.2 mm TPU mesh through which the CO_2_ needle can be inserted. A fine fly egg collection net is also heat-bonded into the cap during printing: after the first ten 0.12 mm layers, printing is paused to place and secure the net over the unfinished part, then resumed to fuse it into the print. The net prevents flies and external parasites such as mites from entering the vial while allowing air and CO_2_ to pass. The vial cap is modified for the backup vial to which the robot returns individual flies that need to be stored. A small incision is made on the top TPU mesh, while the fine mesh is omitted. The glass pipette can be inserted through the incision and the mesh flexibly wraps around the pipette. This prevents flies in the vial from escaping while new fly is being added. Once the pipette retracts from the vial, the incision flexibly closes. Standard cellulose and foam fly stoppers were tried first but deform after initial use and could not be handled repeatably by the robot.

#### Single fly housing plates and shelves

A plate with two 32 square-shaped wells on each side is printed in PLA (Supplementary figure 4A-D). The slicer settings omit the shell of the bottom wall and only the 0.7 grid infill is printed such that the floor of the plate is a mesh. This way the plate can be placed on solidified fly culture media providing access to food and water to flies during long-term storage (Supplementary video 2). Additionally, the net prevents the flies from sticking in food and facilitates CO_2_ flow during anaesthesia when the flies need to be extracted from the wells. The rest of the plate is printed with 100% infill and 0.12 mm line height. There is an impression at the middle of the wider side of the plate that fits the geometry of the robots gripper finger. The plate is covered by laser-cut 1 mm thick acrylic lid. The lid has 64 1 mm holes that, when the lid covers the plate, they align with the centre of each square well (Supplementary figure 4C). A round protrusion at the centre of the plate facilitates vacuum gripping of the plate with a suction cup, and holds the acrylic lid in place. Taping over the protrusion can also be used to keep the lid secure when a person is handling the plate (Supplementary figure 4D). One 64-well plate is affixed to the bench top to serve as the lid holder.

#### Plate storage shelves

The shelving for single fly housing plate storage was laser cut from 3 and 5 mm transparent acrylic sheet and assembled to form a 2 x 10-slot shelving system (Supplementary figure 4B, 4E). The connections are glued together, the two shelves are connected with M6 screws and affixed stably to the bench top using aluminium profiles.

#### 3D print models

All models for 3D printed components were designed in AutoCAD or TinkerCAD and can be found on the GitHub repository. Unless stated otherwise, all models are sliced in Cura, with 100% infill and 0.12 mm line height, and printed in black matte Ultimaker PLA on Ultimaker 5S printer with a glass print bed and standard Magigoo print adhesive. The components that need to be affixed to the benchtop also have M6 screw fittings 25 mm apart that match the benchtop M6 tap raster.

### Coordinate frames

Once all bench top components are installed on the bench top, their pose and coordinates are input in the robot’s wobj variable. This procedure must be performed once when the cell is first installed or if changes are made to the composition of the workbench. The pose of each component in the cell was taught manually using the robot’s lead-through function. A local coordinate frame was defined for each component using three reference points: the first two points define the positive direction of the x-axis and the third point defines the y axis (Supplementary figure 6). The origin of the right-handed Cartesian coordinate system is calculated from the first three points (Supplementary figure 6). These points were recorded in the robot base frame using a pointed calibration probe (an extended aluminium cone mounted on the end effector arm) with its tip defining the tool centre point (TCP). The resulting transformation related the local component’s frame to the robot’s base frame. The complete registration procedure was repeated at least three times, yielding a positional repeatability of 1 mm.

### Behaviour platform

The robot cell is organized around a multiplexed behaviour platform we previously developed for manual phenotyping of learning behaviour in flies and the circuit design and hardware components can be found in^1^. Before integrating the platform into the robotic cell we performed two design changes compared to the previous version where because we saw an opportunity for optimization of the assay itself^1^. First, the 64 electromagnetic relays previously used to gate individual shock delivery were replaced with 16 four-channel MOSFET optocouplers (Vishay KP847PH). This improved switching response from 200 ms to 200 ns, sharpening the timing of closed-loop shock delivery for operant conditioning assays, and it reduced the control PCB footprint to 100 x 80 mm^2^. Compared to the previous design of the platform, there was also a reduction in maximum voltage rating from 230 V to 70 V, which was immaterial at the stimulus amplitudes usually used in the assay (30 V).

Second, the LCD panel (LG 25BL56WY, 25”, 1920 × 1200) was stripped of its casing and its power delivery, LCD control, and backlight circuitry was relocated to the workbench at a distance from the panel. Thermal pads (Iceberg Thermal DRIFTIce, 2 mm, 13 W/mK) are applied to the back of the LED backlight for better heat dissipation. The LCD panel is also elevated for airflow to 125 mm on mounting posts (Thorlabs P100/M, 100 mm height) holding 3D printed right-angle brackets. This modification was done primarily to reduce the screen surface temperature from 28.3 *±* 2*^↑^*C to 26.5 *±* 0.1*^↑^*C. These changes also allowed the LCD panel to be rigidly fixed within the robot’s reachable workspace. Freeing space was beneficial for both human and robot handling, as it allowed more flexible and comfortable arm trajectories.

### Image acquisition and fly detection

After the robot distributed the anaesthetised flies on the CO_2_ pad, images were acquired with a Canon EOS 2000D DSLR camera (6000 x 4000 pixels) and a 70 mm lens at f/8, ISO 100, and 0.25 s shutter speed. Image processing was implemented in Python using OpenCV (cv2) and NumPy. Four ArUco markers (dictionary DICT_6X6_100; IDs 1 - 4) were used to locate a predefined 120 x 120 mm region of interest (ROI) on the pad. When all four markers were detected, their corners were used to calculate a perspective transform; the ROI was resampled at 24 pixels/mm (2880 x 2880 pixels) and rotated 90° clockwise. The ROI origin was defined in the robot firmware, allowing detected fly positions to be converted to robot coordinates.

Fly detection used two stages. First, OpenCV’s blob detector identified candidate contours using threshold levels from 0 to 255. Candidates were filtered by area (500 - 4000 pixels), circularity (minimum 0.1), convexity (0.5 - 1.0), and inertia ratio (minimum 0.15). In the second stage, the image was converted to HSV and thresholded over the range (2, 162, 0) to (30, 255, 255). The resulting mask was dilated three times and eroded once, then inverted before blob detection. These candidate blobs were filtered by area (800 - 1800 pixels), circularity (0.01 - 1.0), convexity (0.01 - 1.0), and inertia ratio (minimum 0.01). Candidate contours from the first stage without an overlapping bounding box from the second stage were treated as uncategorised and excluded. The algorithm retained flies only if they were still far enough apart for individual robotic pickup; overlapping flies and objects that failed the filters were excluded. Detection parameters are sensitive to illumination and were tuned to the room’s lighting conditions during cell installation.

### Sex classification and coordinate transfer

For each fly detected in the second stage of fly detection, a rotated crop was extracted using the contour’s minimum-area bounding rectangle. To enhance posterior abdominal pigmentation, the blue-channel intensity was subtracted from the red-channel intensity. The difference image was thresholded at 32 (binary threshold), and blobs were detected using area (1 - 500 pixels) and convexity (minimum 0.1) filters. A detected blob classified the fly as male; flies without a detected blob were classified as female.

Fly positions were calculated as the centre of each detected contour’s bounding rectangle and recorded with the sex classification. The detection and classification algorithm was evaluated using an annotated image dataset available for download from GitHub repository. Detected positions were converted from image coordinates to coordinates in the robot’s workspace and sent to the robot for subsequent handling.

### Software and control architecture

The system is coordinated by five Python programs running on a Windows computer and a Raspberry Pi, together with robot routines written in ABB RAPID, ABB’s proprietary high-level programming language for industrial robot control. The software was originally developed in Python 3.9 and was maintained through Python 3.12, the version used when the experiments for this manuscript were conducted. The packages used are well-established open-source projects with active community support. They remain available and maintained for current Python releases, including Python 3.14, the latest stable release series at the time of writing (Python.org). Dependencies are listed in `requirements.txt` and can be installed with `pip install -r requirements.txt`. The software, including the RAPID code, will be made open source upon publication to support reuse and further development.

On the computer station (Windows), `master_control.py` monitors the Raspberry Pi’s FTP server for `start_experiment.rdy`, which the robot uploads after preparing the flies. When the flag appears, the program runs the conditioning experiment, deletes the flag, and `uploads experiment_done.rdy` to signal the robot to continue. This cycle repeats until the programmed number of experiments is complete. The program send_flies_coordinates_to_yumi.py monitors for `robReadyForCO2Camera.rdy`, uploaded by the robot when fly coordinates are needed. It triggers image acquisition, processes the image to determine fly positions and sexes, uploads the resulting `flies_coordinates.csv file`, and deletes the request flag. The third computer program, `keep_me_up.py`, runs in the background and periodically adjusts the system volume to prevent the computer from sleeping. On the Raspberry Pi, server.py creates and maintains the FTP server used for communication between the computer and robot. The program `flyGridMove.py` monitors for `move_grid.rdy` and activates the vibrating sieve while the flag is present. The behaviour platform controls CO_2_ delivery through an Arduino-controlled solenoid. The robot’s movement, coordination, and communication routines are programmed in RAPID. Together, these components coordinate the workflow until the programmed number of experiment cycles is complete or the supply of fly vials is exhausted.

### Drosophila stock and rearing

Same rearing protocol was used as in^1^. Briefly, flies were reared on cornmeal agar medium containing 6.2 g agar, 58.8 g cornmeal, 58.8 g brewer’s yeast, 0.1 L Migros M-Classic grape juice, 4.9 mL propionic acid, 26.5 mL Moldex methylparaben, and 1 L Lausanne tap water. All genotypes were obtained from the Bloomington Drosophila Stock Center (BDSC) in 2020. F0 stocks were maintained on the same medium at 21 ± 1°C without controlled light cycles and contained 20–60 flies per vial, without larval-density control.

The maternal F1 experimental generation was established in three replicate vials using two sets of F0 flies from stock vials. Nine females and five males were housed together per setup and allowed to lay eggs for three days before removal. F1 progeny were collected 12 days after the final day of egg laying. Experimental F2 vials were established identically from each F1 replicate. Nine days after removal of the F1 adults, vials were checked daily for eclosion. The first emerging flies were discarded, and flies emerging during the following day’s peak were collected. Approximately 40 males and 40 females were transferred to a fresh-medium vial and assayed at five days of adult age (5 days post-eclosion). Experimental generations were maintained at 26 ± 0.5°C and 50 ± 5% relative humidity under 12-h light/dark cycles with instantaneous transitions. In benchmarking comparisons between human and robot handling flies from the same vials were split and assigned randomly to each handling mode.

### Operant conditioning assays

We used an associative visual operant place-learning paradigm, in which unrestrained flies explored a Y-maze and make sequential binary decisions at the junction of its three arms. Two arms of the maze are illuminated with blue colour while the third one is green. Following an initial 5-minute measurement of unconditioned, innate biases, conditioning was initiated such that each decision could result in the introduction, continuation, or removal of foot shock. For each genotype, two biological replicates derived from independent sets of parents and reared in separate vials were tested. Each replicate comprised 32 flies, equally divided by sex and between conditioned and control groups. Conditioning was performed under the same environmental conditions used for rearing (26 ± 0.5°C and 50 ± 5% RH). Flies were lightly anaesthetised using CO_2_ (10 L/min flow through the CO_2_ pad) and loaded into the Y-maze arenas with an aspirator in the case of manual handling and with the vacuum pipette tip with robotic handling. When conditioned, the flies received a shock upon entering the green arm, which ceased when they exited it, combining positive punishment with negative reinforcement.

### Definitions of behavioural measures

Behaviour measures are defined as in^1^. All behaviours are measured first during the first 5 minutes when the flies are not shocked and behave spontaneously (“baseline” measurement, “innate” behaviour) and then during 20 minutes when they are shocked (“conditioned”) or not shocked (“control”). All behaviours are measured for individual flies. Mean behaviour is calculated as average behaviour of all individuals of the same genotype.

**Task performance or colour preference** of a fly is the fraction of time a fly has spent in the green arm within a time-frame. **Handedness or turn bias** of a fly is the absolute value of the total number of right turns subtracted from the total number of left turns, divided by the total number of turns taken by the fly at any given time frame. **The learning score** measures the change in colour preference over time, and it is calculated by dividing each experiment into ten 2-minute intervals and taking the negative mean of two Spearman’s rank correlation coefficients: (1) the correlation between interval index *i* and the proportion of time spent in the shock arm during that interval, and (2) the correlation between interval index *i* and the proportion of arm visits directed toward the shock arm during that interval. **Activity** is the average velocity of the fly within a timeframe.

### Data filtering

We applied two levels of data quality filtering before behaviour analysis. For retention estimates, we used a **lenient retention criteria** intended to identify flies that remained minimally active after handling and setup: flies were required to have a mean activity greater than 0.1 mm/s and to make at least three choices. We allowed small numerical tolerances when evaluating time fractions and bias measures to account for noise in the fly tracking algorithm. The assay tracking algorithm returned missing values (NAs) when a maze was empty or when it detected more than one blob, corresponding to multiple flies, within a maze. We excluded these invalid mazes from behavioural analyses. The lenient criteria generally reflects dead or severely injured flies, an important metric for assessing the physical impact of robotic handling and the system’s ability to minimize handling-induced injury.

However, for most behavioural analyses, especially learning, it is necessary to apply a **stricter behavioural filtering criteria** that excludes flies that are still active, but exhibit strong outlier behaviour indicative of injury (Supplementary table 2). Individual flies were excluded if they died in the maze or were inactive as per lenient criteria, but also if they never entered the shocked arm/colour, spent at least 90% of the assay in the shocked arm/colour, or made fewer than eight choices during conditioning. We additionally required activity during conditioning period to be greater than 0.2 mm/s. Over the 1200 s conditioning period, this threshold corresponds to 24 cm of movement. At approximately 3 cm per choice, that amount of locomotor activity results in about eight choices. Requiring at least eight choices also ensured a minimum amount of behavioural data necessary for estimating learning performance across the ten 2 minute bins (see^1^ for learning score calculation), which otherwise returns NA as measurement.

Together, these criteria removed extreme or absent engagement with the assay, which can arise when a fly is injured during loading, it is sick or otherwise unwell, or if it presents outlier preference phenotypes, such as shock preference, which can be confounded with injury. These criteria were applied to both conditioned and control flies, where applicable.

### Measures of variability

The extent of behavioural individuality within a group of flies was quantified as the differential entropy of the behaviour’s probability density function, estimated over its theoretical range (H) as described in^1^. For Gaussian distributions, entropy scales with the logarithm of the standard deviation^46^ and is therefore consistent with conventional measures of variability.

## Statistical analysis

All statistical tests are performed in R environment (version 4.0.2 or higher).

## Data availability

Behaviour data will be made publicly accessible upon publication in the Fly Robot Cell gitHub repository. Raw timing logs are recorded in the supplementary file timing_data_robot_manual.csv in tidy format.

## Acknowledgements

We would like to thank Bloomington Drosophila Stock Center (funded by NIH funding P40 OD018537) for providing Drosophila stocks. This study was funded by the ELISIR scholarship.

## Author contributions statement

R.M., T.Š., I.T. and A.M.J. conceptualised the study and designed the robot cell. R.M. and T.Š. wrote the software. R.M., I.T. and A.M.J. designed and built the cell hardware. R.M. prepared and performed the fly experiments, analysed the data, and edited the initial manuscript draft. R.M. and A.M.J. interpreted the data. R.M., T.Š., I.T. and A.M.J. read and edited the manuscript. A.M.J. wrote the manuscript, secured funding and supervised the study.

