## Supplementary figures, videos and tables for "End-to-end automation of behavioural experiments with a collaborative robot"

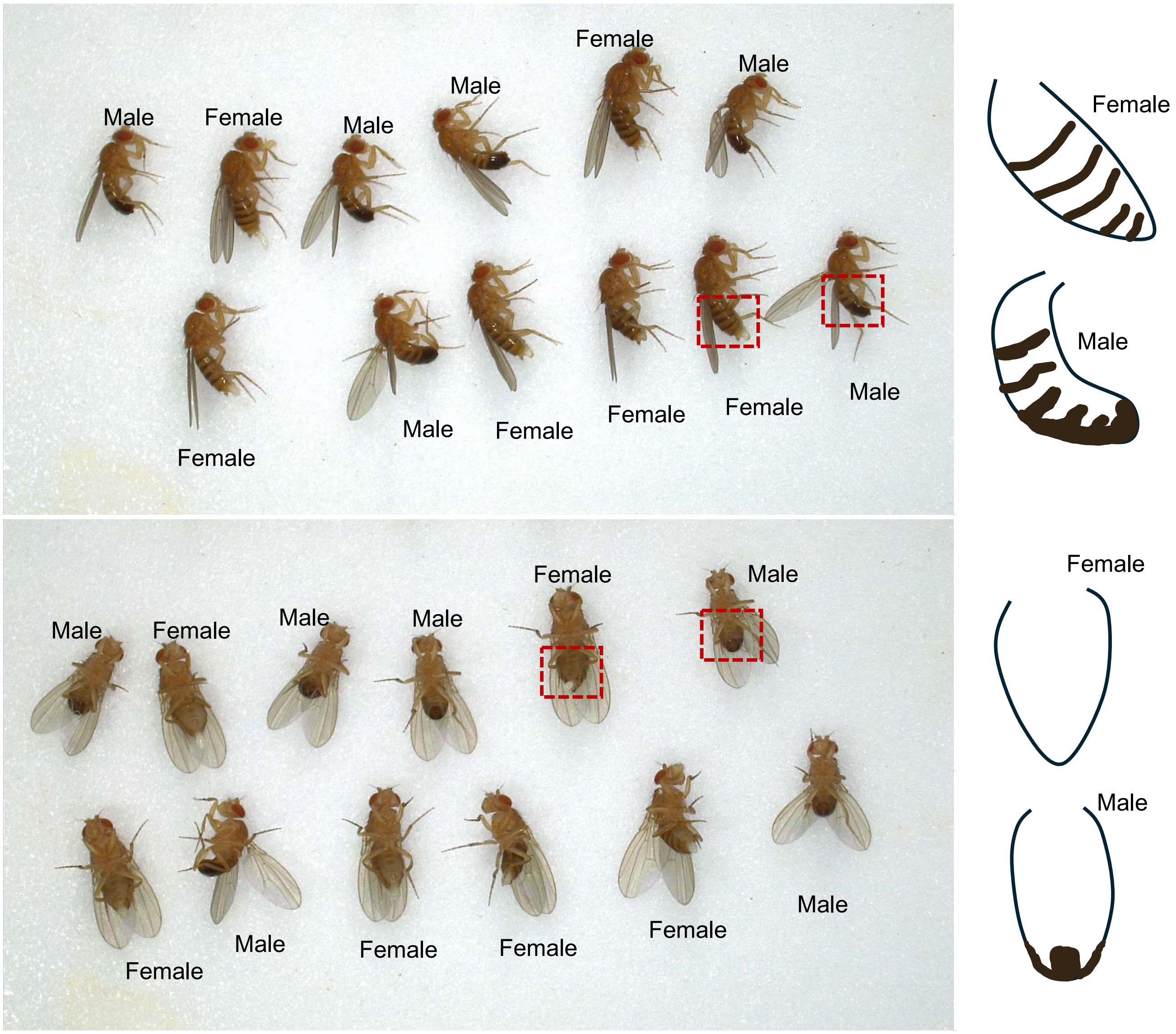


**Supplementary Figure 1. Fly sex determination.** Male and female pigmentation and body morphology differ, particularly in the abdomen (red box). Males have a much more curved abdomen, with a more rounded posterior end and a pronounced pigmentation as exemplified by the abdomen sketches on the right. These features are used in detecting males and females by both humans and the robot.

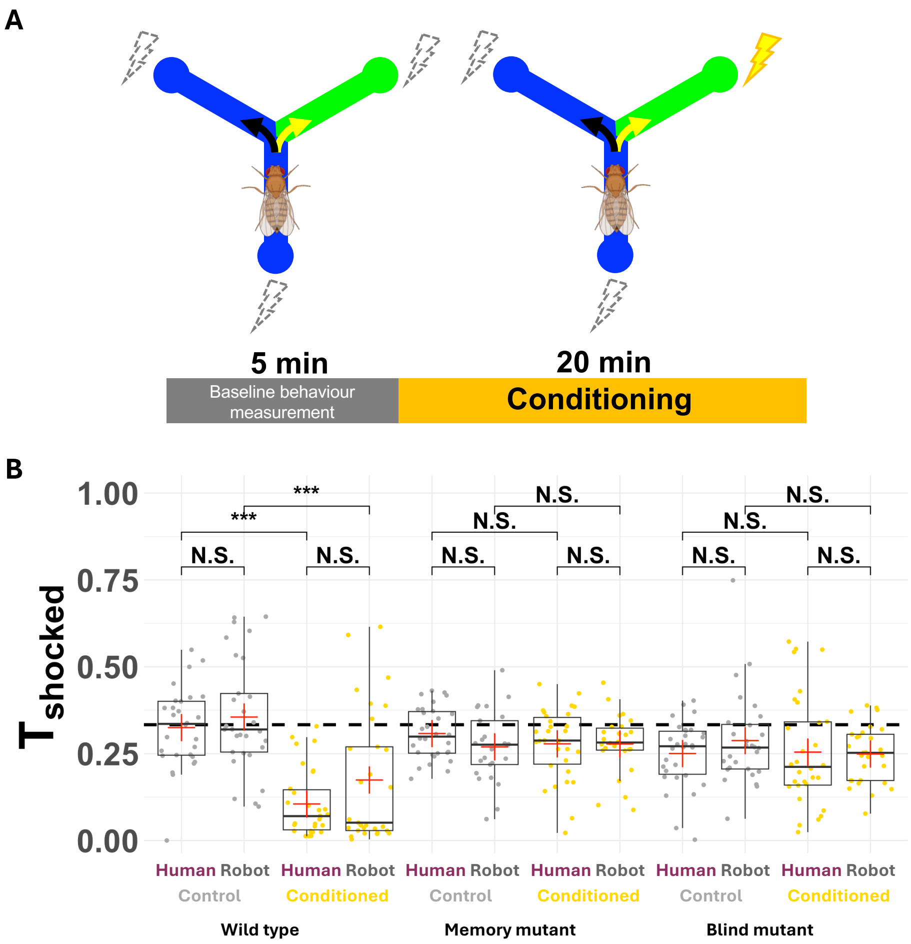


**Supplementary Figure 2. Conditioning paradigm and biological validation. A)** Operant conditioning learning paradigm implemented in this study. In control experiment, the baseline measurement (without shock) extends throughout the entire 25-min assay. **B)** Task performance ($T_{shocked}$ , proportion of time on green colour) in green place learning task (green arm is shocked) in the mutant behaviour experiment. Three genotypes were used in this experiment, a wild-type genotype (DGRP-362) compared to the memory mutant (*dnc*$^{1}$) and the blind mutant (*norpA*$^{EE5}$). 'Robot' (grey) and 'Human' (magenta) represents automated and manual handling respectively. Grey and yellow points represents control and conditioned flies. Wild type flies learn to avoid the green arm and the mutants do not, in both types of handling. There is no significant difference in task performance between automated and manual handled flies for corresponding groups of flies (N = 384, n = 32 vs 32 for the t-test).

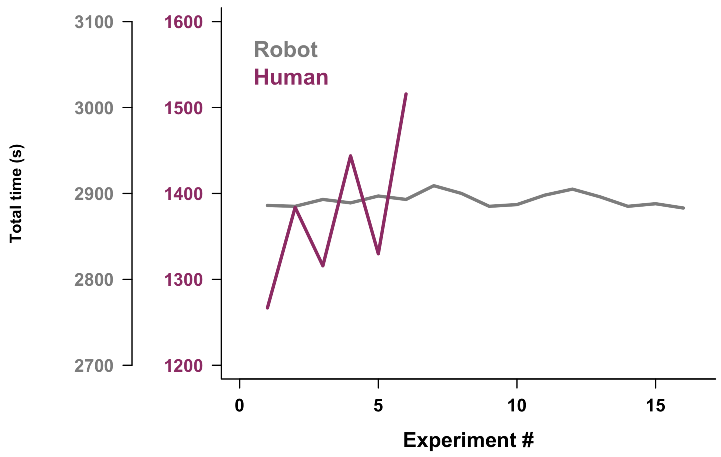


**Supplementary Figure 3. Time invested per experiment.** Invested time per experiment for the robot (setup including fly sexing + serial dismantling time, but excluding assay time when robot is idle) is shown in grey while human (average fly sexing + setup + group dismantling time, but excluding assay time) in magenta. Note that in the human-conducted experiment, males and females are separated two days prior to the experiment and the dismantling time is not serial.

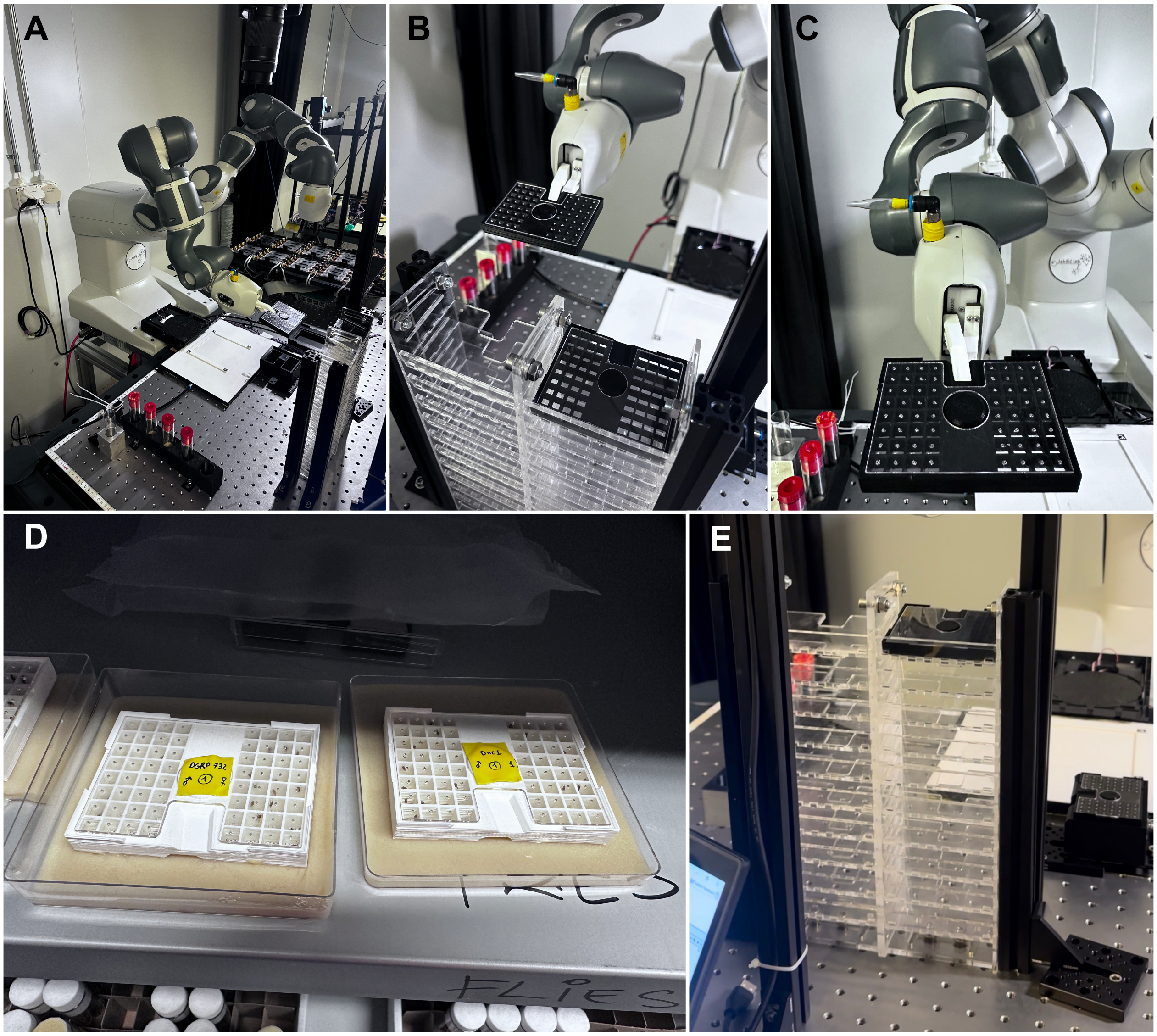


**Supplementary Figure 4. Long-term single fly storage.** The flies can be picked from the $CO_2$ pad or from the Y-maze and placed in the 64- well plate for single-fly housing. **A-C)** The robot handles the plates using the gripper indentation that fits its fingers. **D)** The plates can also be handled the same way by human experimenter and placed on fly culture media for long term storage and maintenance of live flies. **E)** The robot can store, track or access plates in the shelving system that can hold 20 plates. Because humans cannot move linearly as steadily as the robot, or track and memorise the plates positionally, it is advised to affix the plate lid with a tape and label each plate immediately after removing from shelving, to avoid accidentally opening the plate.

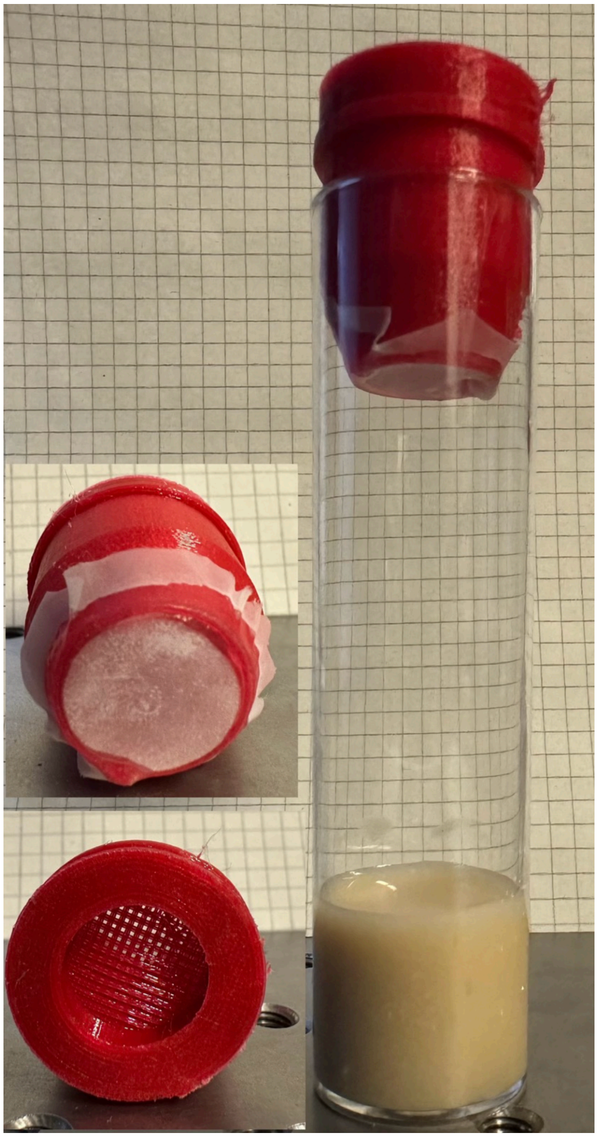

**Supplementary Figure 5. Custom Drosophila vial cap.** The cap was designed to fit a standard Drosophila culture vial (25 mm diameter). The cap is hollow, but closed at the top and the bottom. The inset top photo shows a close-up of the fine mesh on the bottom that prevents mites from entering the vial and flies from entering the cap. Without the net, the flies trapped in the cap following anaesthetized could not be transferred onto the $CO_2$ pad in a controlled manner. The bottom inset photo shows the 3D printed net that allows insertion of the $CO_2$ needle into the closed vial.

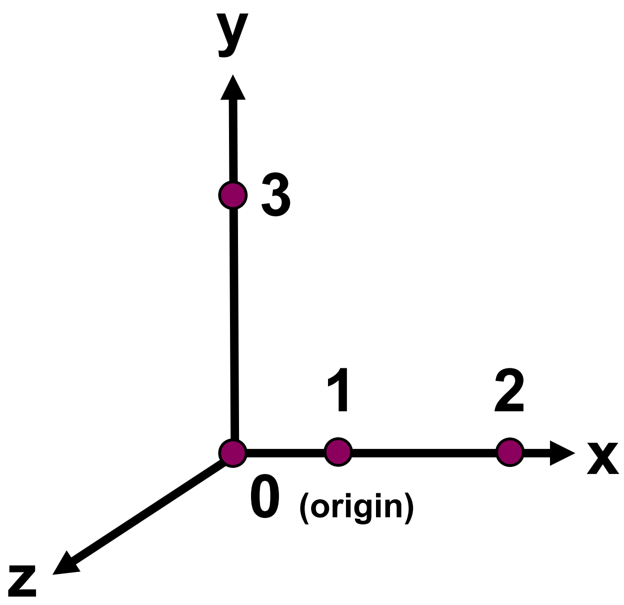


**Supplementary Figure 6. Coordinate frame.** Points 1-3 are used to determine the origin (0) of the local right-handed Cartesian coordinate frame for each workspace object.

#### Supplementary videos

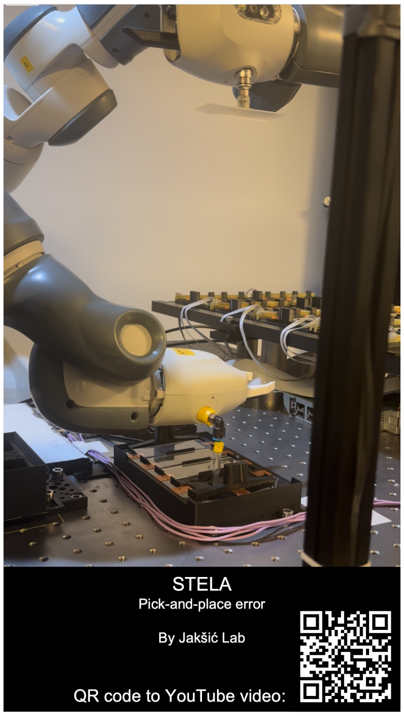


**Supplementary Video 1. Pick-and-place error.** Male and female pigmentation and body shape differ. Males express a much more curved abdomen, a more rounded posterior end with a pronounced pigmentation as exemplified by the abdomen sketches on the right. These features are used in detecting males and females by both humans and the robot.

### Supplementary tables

**Supplementary Table 1. Experiments and genotypes used for different performance measurements.** The robot's performance was tested within the scope of two experiments that used different mutant genotypes (A.) and wild-type genotypes (B.) listed in the table. Coloured cells indicate which genotypes were handled by the robot (grey), by the human (magenta) and by both (black). Only the genotypes that were handled by both the human and the robot were used to compare biological outcomes. The mutant behaviour experiment was used to measure only the biological outcomes. The cycle performance was measured only within the wild-type behaviour experiment (blue).

| Genotype | Expected phenotype | Genotypes handled by robot | Genotypes handled by human | Overlap for comparing biological outcomes | Used to measure quality filtering and cycle time |
|---|---|---|---|---|---|
| **A. Mutant behaviour experiment** | | | | | |
| DGRP362 | Wild-type, good visual learning | | | | |
| *norpA*$^{EE5}$ | Blind, deficient learning | | | | |
| *dnc*$^{1}$ | No memory, deficient learning | | | | |
| **B. Wild-type behaviour experiment** | | | | | |
| DGRP 181 | Wild-type | | | | |
| DGRP 348 | Wild-type | | | | |
| DGRP 358 | Wild-type | | | | |
| DGRP 361 | Wild-type | | | | |
| DGRP 57 | Wild-type | | | | |
| DGRP 85 | Wild-type | | | | |
| DGRP 149 | Wild-type | | | | |
| DGRP 176 | Wild-type | | | | |
| DGRP 229 | Wild-type | | | | |
| DGRP 336 | Wild-type | | | | |
| DGRP 352 | Wild-type | | | | |
| DGRP 365 | Wild-type | | | | |
| DGRP 41 | Wild-type | | | | |
| DGRP 45 | Wild-type | | | | |
| DGRP 48 | Wild-type | | | | |
| DGRP 88 | Wild-type | | | | |

**Supplementary Table 2. Behaviour data filtering before cycle benchmarking.** To benchmark robot's against human's performance two types of filters were applied to the data from the 24-h wild-type behaviour experiment. Lenient filtering removes dead and severely injured flies, while stringent filter removes those and the flies that show strong outlier behaviours that may bias the complete behaviour scoring. Behaviour measurements are then calculated for all flies that pass this filter. Retained flies for which any behaviour measurement returns NA are also removed from the analysis, leaving the final list of scored flies (n).

| | Lenient activity filtering | | | Stringent behaviour filtering | | |
|---|---|---|---|---|---|---|
| | Robot Subset 6 wild-type genotypes | Robot All 16 wild-type genotypes | Human 6 wild-type genotypes | Robot Subset 6 wild-type genotypes | Robot All 16 wild-type genotypes | Human 6 wild-type genotypes |
| **Total number of handled flies (N)** | 384 | 1024 | 336 | 384 | 1024 | 336 |
| **Flies retained after filtering** | 350 | 961 | 280 | 266 | 764 | 180 |
| **Scored flies (n)** | 350 | 961 | 280 | 266 | 761 | 179 |
| **% flies fit for analysis (n/N x 100)** | **91.15** | **93.85** | **83.33** | **69.27** | **74.32** | **53.27** |
